# Coarse-graining reveals collective predictive information in a sensory population

**DOI:** 10.1101/2025.10.18.683195

**Authors:** Adam G. Kline, Maciej Koch-Janusz, Aleksandra M. Walczak, Thierry Mora, Stephanie E. Palmer

## Abstract

Biological systems perform complex computations using hundreds of individual actors, but they do so efficiently and in a way that can be read out and interpreted by other biological networks. Coarsegraining may allow for key collective features to be effectively and efficiently communicated. In the brain, early sensory systems perform prediction, which can compensate for lags in neural processing. This computation is collective, meaning it relies upon interactions between many neurons, and operates in complex, dynamic natural environments. Taking these two facets of biological complexity together, we search for maximally-predictive collective variables in large groups of retinal ganglion cells responding to dynamic natural visual scenes. To find collective variables that best capture predictive computations in the neurons, we apply a tractable, approximate implementation of the information bottleneck method to neural data. We infer a lower-dimensional representation that is maximally informative about the future neural activity. We observe scaling relationships between this mutual information estimate, neural subset size, and information decay timescale. Further, the structure of collective modes changes for predicting at short versus longer timescales.

## INTRODUCTION

Prediction is ubiquitous in biology, allowing organisms to make use of correlations in their environment and overcome processing lags. The success of a species depends on its ability to survive and reproduce, and prediction is crucial for reacting quickly in life-or-death situations. Prediction has therefore been explored as a normative principle for understanding how organisms assign utility to various components of incoming signals in early stages of sensory processing [1–4]. In the vertebrate visual stream, signals are propagated through several layers of neurons, each with significant sensory delays, that prediction may help ameliorate. This begins in the eye, which can optimally predict visual signals in both artificial and naturalistic visual environments [5–10]. While optimal prediction has been observed for small groups of cells in the vertebrate retina under simple stimuli [7], we expect that this computation should not only extend to richer stimuli, but should also continue along the visual stream in order to produce the fast reaction times in natural scenarios [3, 11–13].

Visual signals are first processed in the retina, then sent downstream to ultimately drive behavior. While neurons in the retina encode all that the brain sees, the visual information they carry must be decoded into signals or features which are useful to the organism. Abstractly, this decoding problem is difficult because the retinal output is a high-dimensional random variable consisting of the collective state of all ganglion cells. On the other hand, this output contains correlations, both because they are present in the stimulus [14–22] and because there are interactions between neurons [23–28]. These correlations reduce the space of likely codewords and may therefore enable decoding. In particular, given the importance of fast prediction to survival, correlations in the retinal output should allow for neurons downstream to read out signals which are informative of the future. To address this question, we search for predictive coarse-grained features of retinal responses to naturalistic stimuli and explore the correlation structure that allows for this predictability.

To study how the collective properties of retinal responses might allow for downstream prediction, we analyze correlations in the simultaneous activity of a large number of neurons stimulated by a range of naturalistic images. Given the wide range of timescales present in natural visual environments [18] and the retina’s ability to resolve very fast features [29, 30], we focus on long recordings with high temporal resolution.

To find and study predictive features in the retinal population code, we apply recent machine learning methods which leverage concepts from information theory [31–34]. At its core, our method solves a variational version of the predictive information bottleneck problem [35] by simultaneously searching over predictive coarse-grained features and performing inference. By using the expressive power of neural networks to solve an inference problem, this method is able to estimate information quantities on a state space of unprecedented size, that is, the state space of joint activity among hundreds of neurons.

In doing so, we see how many linear features are needed to be predictive, providing support for the feasibility of compression-based prediction in the retina and more broadly neural coding.

## RESULTS

### A. Recordings of retinal neuron population activity

In order to investigate the statistics of retinal outputs, we examine response data taken from 93 retinal ganglion cells (RGCs) in the salamander retina under naturalistic stimuli [36], as described in [10, 37]. The data collection process is outlined in Fig. 1. A retina taken from a larval tiger salamander was placed on an electrode array with a density of roughly one electrode per RGC. These data were then spike-sorted, yielding the times at which each neuron fired an action potential. A binary representation of the neural code was generated by discretizing time into time bins of size Δ*t* = 1*/*60 s, and assigning the state “1” to a neuron in any time bin in which it fired a spike and a “0” otherwise. In our data, the marginal probability of the “1” state across all neurons and time bins can be as high as 0.016 and as low as 0.003, depending on the stimulus.

**FIG. 1.**
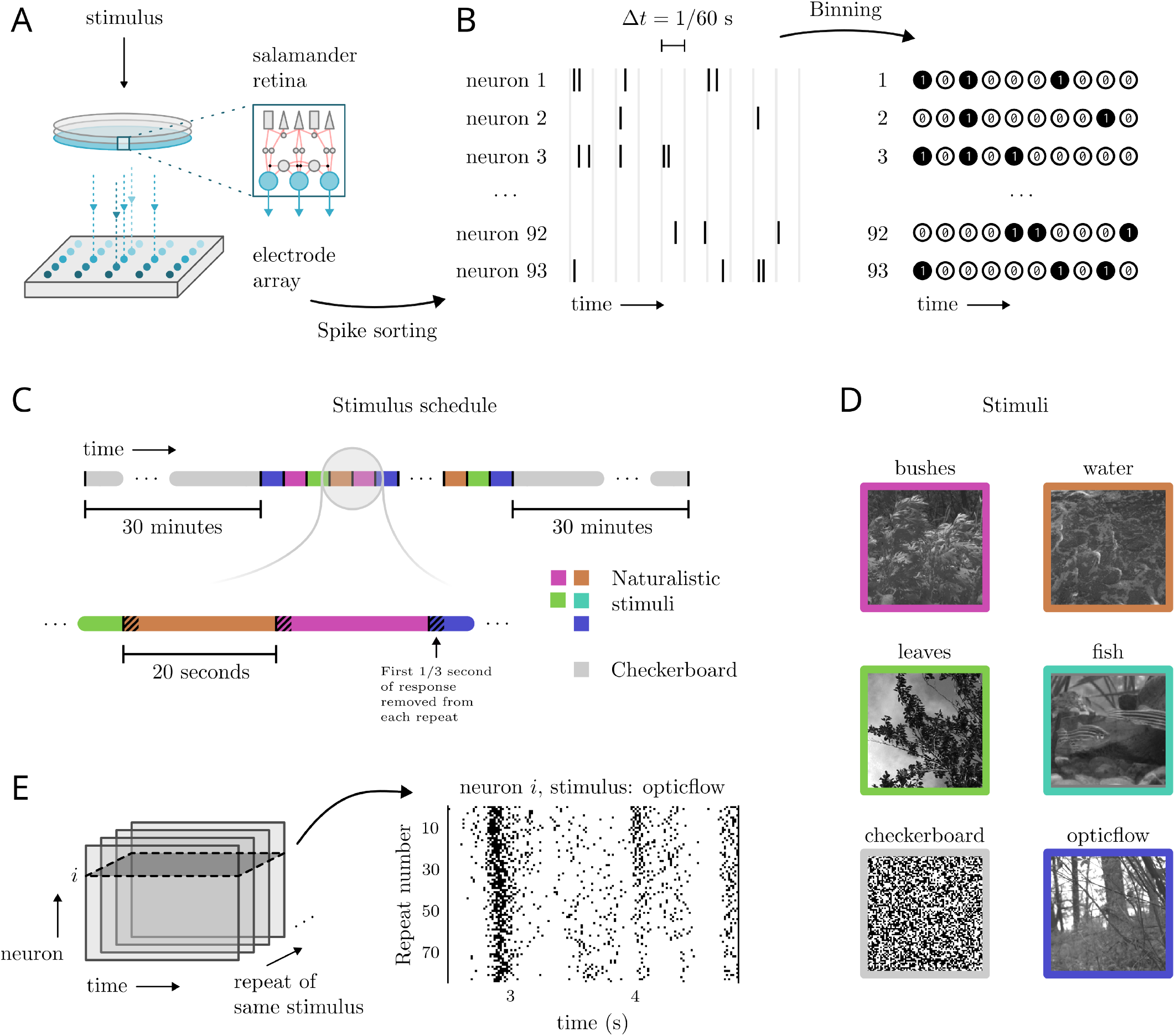
Neural data. Experimental protocol enables simultaneous recording of responses across a large population of salamander retinal ganglion cells under repeated, naturalistic stimuli. **A**. Stimuli are shown to a salamander retina while ganglion cell responses are recorded by an electrode array. **B**. Following spike sorting from electrode array, neuron spike times are binned into Δ*t* = 1/60 second time bins (aligned with stimulus movie frames), which yields a binary representation of the neural code. **C**. Naturalistic stimuli are shown at 60 frames per second to the retina in random order, in 20 second intervals. In our analysis, the responses from the first 20 frames (0.3 seconds) of each interval are removed, as they contain transients due to switching stimulus. At the beginning and end of the experiment are 30 minute intervals of checkerboard stimulus. **D**. Snapshots taken from each stimulus. **E**. Each stimulus is shown to the retina ∼ 80 times, allowing us to observe noise correlation effects.

Five different naturalistic stimulus clips were used, each 20 seconds long. These were shown to the retina in random order for 141 minutes, yielding roughly 80-90 repetitions of each stimulus over the course of the experiment. This repeated structure allows us to compare neural responses to a given stimulus across trials, revealing both reliable response features and stochastic effects inherent to the physiology of the retina. Correlations in these latter effects are referred to as “noise correlations” and can be studied by considering the ensemble of repeated trials of the same stimulus. The stimuli themselves are greyscale videos at 60Hz, depicting a range of visual scenes that could conceivably be present in the environment of the specimen. Such stimuli are characterized by heavy-tailed distributions of contrast, velocity, time-, and length-scales, in contrast to simpler, engineered stimuli such as moving gratings [18, 38]. Because the retina encodes its stimuli nonlinearly, an ethologically relevant understanding of these encodings justifies probing responses with naturalistic stimuli [15, 16, 29, 39, 40].

### B. Estimating mutual information

We quantify the amount of information encoded in the outputs of the retina about its future outputs. Generally, given a joint distribution *p*(*x, y*) for two random variables *X* and *Y*, the mutual information between them is

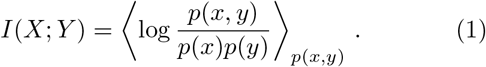

In general, mutual information quantities can be very challenging to compute. A common difficulty is if *X* or *Y* are variables with many dimensions, say *X* ∈ {0, 1} ^*N*^ with *N* large. In such cases, even if given an analytical expression for *p*(*x, y*), the sum in ⟨·⟩_*x,y*_ must typically be approximated. More problematic, when working with data, we must provide an estimate of *p*(*x, y*). This problem is known as inference, and also becomes increasingly difficult as the dimensionality of random variables increases.

In this work, we leverage recently-developed tools to compute a tractable lower bound to (1) [41]. This bound, which relies on an ansatz for *p*(*x, y*) represented using neural networks, is maximized to approach the true distribution. In other words, this method both solves an inference problem and in the process computes an approximation of *I*(*X*; *Y*).

More explicitly, we consider a trial distribution *q*(*x, y*|*θ*) which represents our best guess at *p*(*x, y*), where *θ* are free parameters. The Barber-Agakov (BA) bound on mutual information is given by

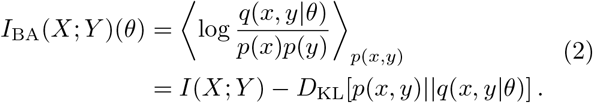

Because the Kullback-Leibler divergence *D*_KL_[*p* || *q*] ≥ 0, we have that *I*_BA_(*X*; *Y*) ≤ *I*(*X*; *Y*) with equality only when *q*(*x, y*|*θ*) = *p*(*x, y*). Therefore, the act of maximizing this bound is, precisely, inference. We parameterize this trial distribution *q* with the ansatz

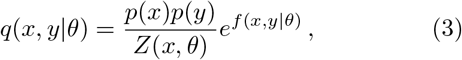

where *p*(*x*) and *p*(*y*) are the true marginals of *x* and *y*, and *f* (*x, y*|*θ*) is computed by a neural network with parameters *θ*. The partition function *Z*(*x, θ*) is then defined by *f* (*x, y*|*θ*) and *p*(*y*) through the requirement that ∑_*y*_ *q*(*x, y*|θ) = *p*(*x*) for all 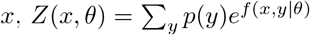.Inserting this ansatz into the BA bound yields the unnormalized Barber-Agakov bound

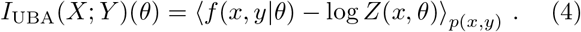

This bound is intractable due to the log partition function. In the noise-contrastive estimate (NCE) bound, a Monte-Carlo estimate of the partition function is computed using minibatches containing *B* independent samples 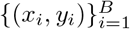from the data, and this effectively reduces variance of the final estimate. Explicitly, the NCE bound is

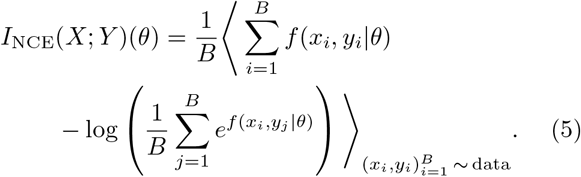

In summary, we aim to compute *I*_NCE_(*X*; *Y*), which is bounded in the following way

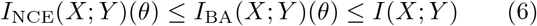

with each bound becoming tighter as *q*(*x, y* |*θ*) approaches *p*(*x, y*). We implement the critic function *f* (*x, y*) using fully connected neural networks *u*_*a*_(*x*) and *v*_*a*_(*y*) with *a* = 1, …, *N*_embed_ as:

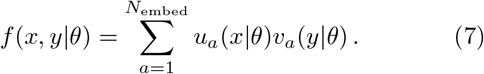

We refer the reader to Refs [32, 41] for a more detailed overview of these bounds.

### C. Predictive information persists at long timescales

The retina needs to reliably encode and transmit visual information for further processing by downstream areas of the brain. Retinal dynamics is shaped by the statistical and dynamical properties of the external inputs. However, downstream neural populations do not have direct access to those properties, other than provided by retinal stream itself. Taking the downstream perspective and looking at the retinal code, itself, as if produced by an autonomous dynamical system, what can we say about its predictability?

We first examine predictive information, defined as the mutual information *I*(*X, Y*) between the joint activity of all *N* = 93 neurons in a single time bin, *X* ∈ {0, 1} ^*N*^, and the same set of degrees of freedom at *τ* into the future, denoted by *Y* ∈ {0, 1} ^*N*^ (Fig. 2A). Our estimate of the predictive information *I*(*X*; *Y*) is given by *I*_NCE_(*X*; *Y*) (in Eq. (5)), evaluated on data which were held out during training of the critic function.

**FIG. 2.**
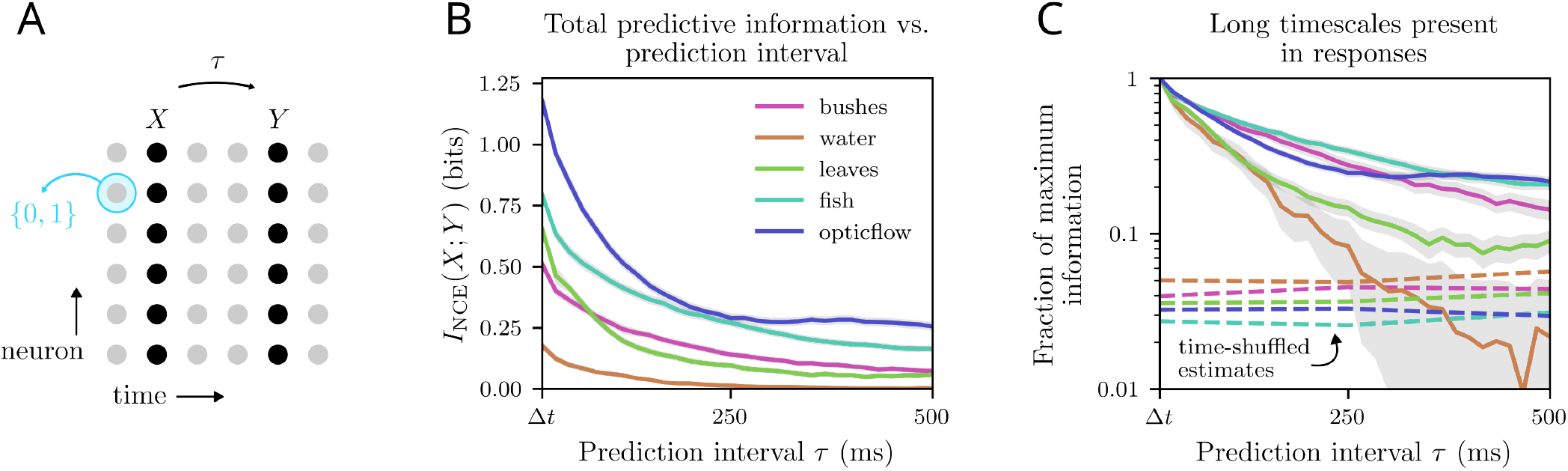
Long predictive timescales under naturalistic stimuli. **A**. We use the NCE bound, Eq. 5, to estimate mutual information *I*(*X*; *Y*) between *X* and *Y*. *X* is the binary codeword of length *N* = 93 representing ganglion cell responses in a single time bin, while *Y* is the codeword *τ* in the future. **B**. Predictive information versus prediction interval for each stimulus. **C**. Predictive information normalized by its maximum value, on a semi-log scale (same color scheme). Each stimulus response displays multiple timescales and correlations out to about 500 ms. Dotted lines depict predictive information estimates after shuffling the time index across repeats. Nonzero quantities below these lines are spurious, as they represent estimates when no information beyond frame index is present.

In Fig. 2B, estimates of *I*_NCE_(*X*; *Y*) are given as a function of the prediction interval *τ*, for each of the 5 stimuli. Since *I*_NCE_ is a lower bound of mutual information, we can conclude that for certain stimuli, such as “fish” and “opticflow”, the true mutual information *I*(*X*; *Y*) achieves large values even at prediction intervals as large as ∼500 ms. In comparison, for the checkerboard stimulus no correlations were observed, up to the refresh rate of the monitor, 30 fps [7]. Additionally, there is variation in the overall scale of information across stimuli, and in the change of shape of the *τ* dependence.

To better observe this variation across natural scenes, we examine in Fig. 2C the decay of predictive information on a log scale, after normalizing by the maximum value of *I*_NCE_(*X, Y*) at *τ* = Δ*t*. These trends show approximately exponential decay for most stimuli over some initial transient period, followed by a plateau. In the “opticflow” stimulus, this profile is pronounced, with the plateau onset occurring after only about 150 ms. One potential worry might be that non-zero estimates of predictive information in this plateau region are due to overfitting. This might occur if the critic function learns to decode frame numbers from present and future frames, which is feasible given that the dataset contains repeats of each stimulus response. To address this, we also provide NCE estimates of predictive information on shuffled data, where this shuffling randomly permutes the time index in a way that is consistent across repeated trials of the same stimulus. If over-fitting due to “memorizing” frame number is occurring, this procedure allows the information estimate to include its contribution while removing contributions from all other correlations in time. Since these shuffle estimates (dashed lines) are below the estimates on intact data, we conclude that long-time predictive information is not likely due to over-fitting by frame memorization. Taking these shuffle estimates as a limit on information, we note that not all of our stimuli elicit responses with any predictability at a time lag of *τ* = 500 ms. Whereas “opticflow” responses contain predictive information at this extreme interval, responses to the “water” stimulus do not, falling below the shuffle threshold at a much earlier interval of 250 ms.

### D. Predictive information is compressible

The substantial mutual information that we find between points separated by large time intervals *τ* indicates that there are features of the neural code that are predictable. Given the importance of prediction for survival, it is reasonable to expect downstream processes to leverage this predictability. However, these processes may not need to perform this prediction task based on the full knowledge of the entire repertoire of retinal activity. For one, we know that the neural code is noisy, with repeated exposure to the same stimulus giving variable responses. Secondly, we know that the optic tectum, the next stage downstream of the retina in the visual stream, has fewer neurons and acts as a bottleneck for the signal. Furthermore, signals which drive motor responses also converge to a small set of motor neurons, indicating a need for compressibility. Such considerations also suggest that an important aspect of retinal computation is its ability to make signals accessible through linear readout downstream [6]. More fundamentally, because of the code’s redundancy, predictability may be supported by an effective representation of lower dimension than the full joint activity. Given a stimulus, how many distinct features of the random variable *X* ∈ {0, 1} ^93^ need to be measured to learn all there is to know about the future state *Y*, and what are these features?

To answer this question, we seek coarse-grained, low-dimension representations *Z* = Λ(*X*) of the joint neural activity *X* in the present that carry maximal information about the future activity *Y*. This approach is the deterministic version of the information bottleneck (IB) for predictive information described in [7], and we follow [32– 34] to find variational solutions using the NCE machinelearning estimate of the mututal information. Suppose *X* ∈ *χ* is the current state of the retinal code and *Y* is the future, as in Fig. 3A. We seek coarse-graining maps Λ : *χ* → ℝ^*K*^ that preserve a maximal amount of mutual information between *Z* = Λ(*X*) and *Y* :

**FIG. 3.**
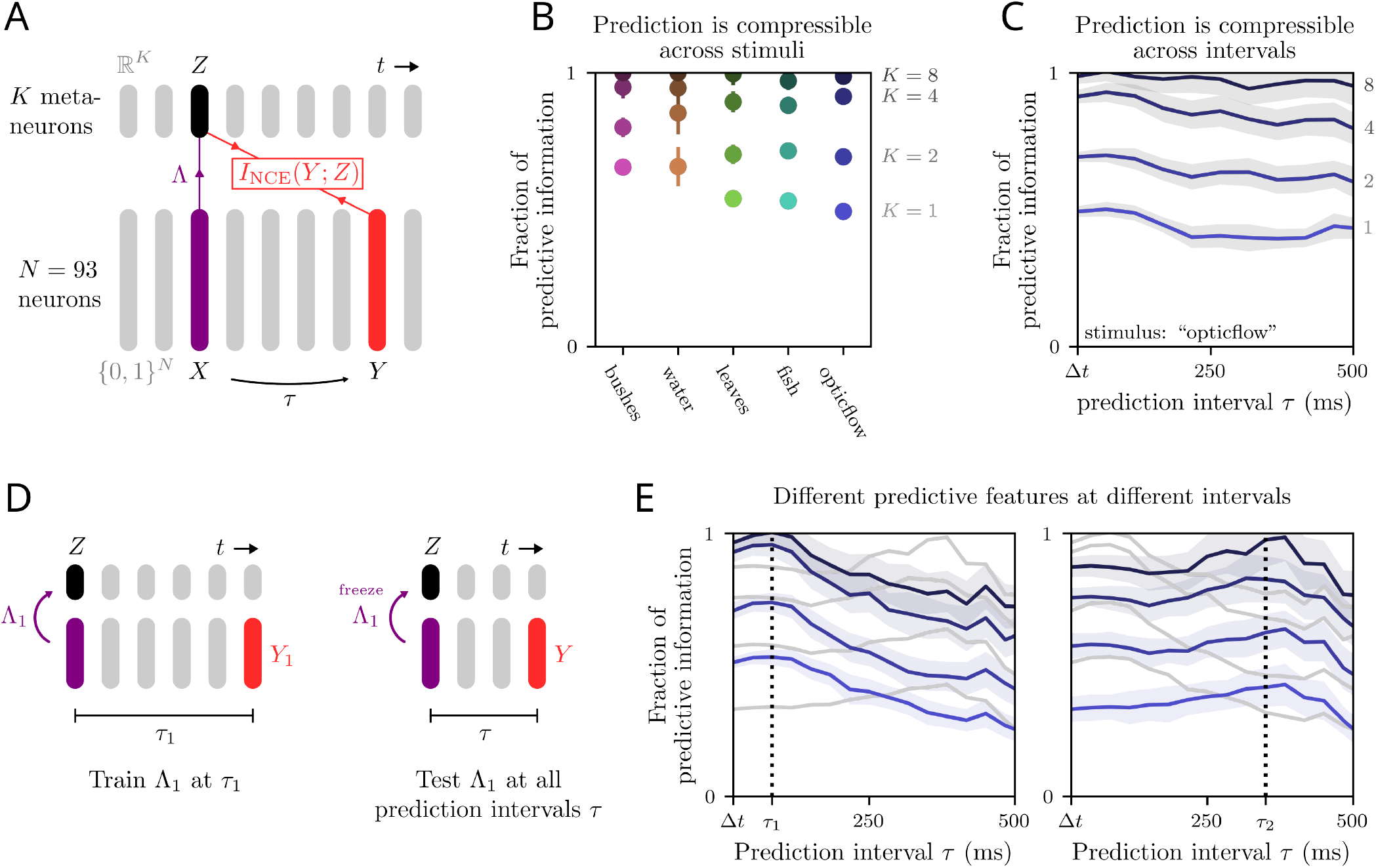
Predictive information is compressible A. Schematic of computation. Joint neural activity *X* is coarse-grained linearly into a compressed, *K*-dimensional representation *Z* = Λ*· X*. The optimal coarse-graining Λ is found by maximizing the NCE lower bound, Eq. 5 along with its variational parameters. **B**. Fraction of total predictive information *I*_NCE_(*Z*; *Y*)*/I*_NCE_(*X*; *Y*) for a single time bin prediction interval *τ* = Δ*t*, for four values of the compressed dimension *K*. **C**. Fraction of total predictive information *I*_NCE_(*Z*; *Y*)*/I*_NCE_(*X*; *Y*) as a function of prediction interval *τ* for the “opticflow” stimulus. Each line represents a different *K* **D**. To test predictive feature generalization, we first find optimal features Λ_1_ at prediction interval *τ*_1_, then measure *I*_NCE_(Λ_1_(*X*); *Y* (*τ*)) for *τ* across the whole range of timescales. **E**. Predictive feature generalization *I*_NCE_(*Z*; *Y*)*/I*_NCE_(*X*; *Y*) for compressed dimensionalities *K* = 1, 2, 4, 8 (same color code as C), at two different training timescales *τ*_1_ = 83 ms and *τ*_2_ = 350 ms, as explained in D.

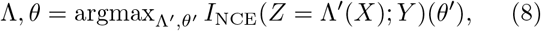

The parameters *θ* for the variational estimate *q*(*z, y θ*) of the true joint distribution *p*(*z, y*) are optimized along with the map Λ. In this work, we restrict our search to linear maps Λ: *Z* = Λ · *X*, where Λ is a *K* |dim(χ)| matrix. We refer to the entries of *Z* as “meta-neurons”, and the row-vectors of Λ as the features that these metaneurons encode. From here on, when the *θ* argument is dropped from *I*_NCE_ that should be taken to mean *I*_NCE_ evaluated after optimizing over *θ*.

Due to the data processing inequality, *I*(*Z*; *Y*) ≤ *I*(*X*; *Y*), where *I*(*X*; *Y*) is the total predictive information. When *K* is increased, *I*(*Z*; *Y*) cannot decrease, and when *K* = dim*χ*, the coarse-graining map Λ becomes invertible, leading to *I*(*Z*; *Y*) = *I*(*X*; *Y*). Together, these constraints tell us that *I*(*Z*; *Y*) is a monotonically increasing function of *K* which saturates at *I*(*X*; *Y*). In particular, to quantify compressibility of the retinal code, we examine how quickly max_Λ_ *I*_NCE_(*Z* = Λ *X*; *Y*), as estimated using the variational method outlined above, approaches its upper bound *I*_NCE_(*X, Y*) as *K* is increased.

We take *X* ∈{0, 1}^*N*^ to be the activity of all neurons in a single time bin and *Y* ∈{ 0, 1}^*N*^ that same activity some time *τ* in the future (Fig 3A) Across stimuli, only about *K* = 8 meta-neurons are required to capture all of the available predictive information for the next time bin, *τ* = Δ*t*, as captured by the fraction of predictive information *I*_NCE_(*Z*; *Y*)*/I*_NCE_(*X*; *Y*) (Fig 3B). Fig. 3C that this result holds for larger predictive horizons *τ*, up to 500 ms. Recall that *N* = 93, meaning the predictive signal can be compressed by more than 90%. However, since each prediction interval *τ* represents a distinctly defined pair (*X, Y*) of input and relevance variables, these 8 predictive directions in state space could be different at different *τ*.

To investigate the dependence of relevant features on prediction intervals, we pick a “training” time interval, for example *τ*_train_ = *τ*_1_ = 83 ms, and solve (8), yielding (Λ_1_, *θ*_1_) (Fig. 3D). The coarse-graining map Λ_1_ represents the *K* linear features of *X* which maximize the mutual information between *Z*_1_ = Λ_1_ *· X* and 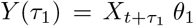 represents part of an estimate of *p*(*z*_1_, *y*). With the features Λ_1_ fixed, we then optimize *I*_NCE_(*Z*_1_; *Y* (*τ*))(*θ*) over *θ* for a range of prediction intervals *τ*. This gives us an estimate of *I*(*Z*_1_; *Y* (*τ*)), or the information between meta-neurons trained to predict at interval *τ*_1_ and the state at all other intervals *τ*. In Fig 3E, we show the dependence of *I*_NCE_(*Z*; *Y* (*τ*))*/I*_NCE_(*X*; *Y* (*τ*)) on *τ* for two different training intervals, *τ*_1_ = 83 ms and *τ*_2_ = 350 ms, and for *K* = 1, 2, 4, and 8 meta-neurons.

For both training intervals *τ*_1_ and *τ*_2_, the predictive features generalize well, but not perfectly, across testing intervals. When the testing interval is equal to the training interval, we see a peak in predictive information carried by these specialized meta-neurons. While it appears to be the case that only 8 meta-neurons trained at any timescale successfully encode the majority of predictive information at all intervals, the two examples we give reveal that these compressions are highly sub-optimal. For example, 8 meta-neurons trained to predict *Y* (*τ*_2_) do worse at predicting *Y* (*τ*_1_) than only 4 meta-neurons trained to predict at that interval. Together, these results suggest that retinal responses for complex naturalistic stimuli encode both general and timescale-specific predictive information, and that these features could be disentangled by downstream neurons solving different prediction problems.

### E. Meta-neurons carry long-timescale predictive information

We have so far shown that retinal responses to naturalistic stimuli exhibit long timescale correlations, and that all of the predictive information can be captured by a few linear coarse-grained variables (meta-neurons). The predictive capabilities of a given set of meta-neurons generalizes to timescales other than the one at which they were trained to predict. Thus, by learning to predict retinal outputs at one specific timescale, neurons downstream of the retina may also learn to predict more generally.

What do meta-neurons encode, and why does this lead to generally good compressed prediction at many intervals? One possible answer is that meta-neurons are encoding slow, long-timescale features. In a theoretical exploration of this idea, Schmitt and coworkers [42] have noted that, in stochastic dynamical systems governed by a time evolution operator *U*,

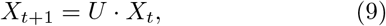

the most informative features identified by predictive IB are given by the eigenfunctions of *U* with the largest eigenvalues, corresponding to the longest timescales. In that setting, each IB feature is associated with a timescale, which can be computed from the eigenvalue decomposition of *U*. Crucially, this identification implies that the compressed features being encoded do not actually depend on the prediction interval, since *U* generates the system’s dynamics. We want to test this idea that meta-neurons (the entries of *Z* = Λ *·X*_*t*_) encode features with long timescales.

We begin by using dynamic mode decomposition (DMD) [43] to construct an estimate of the time evolution operator for the neural dynamics, *X*_*t*_, independently of the IB and its meta-neurons. There is no guarantee that neural activity is well described by a linear evolution operator, in particular because of memory (non-Markovian) effects. To account for non-Markovian dynamics, we operate on a dynamical variable consisting of a time-delay-embedding (TDE) of the neural state: we redefine *X*_*t*_ as the binarized neural responses of all neurons in the 7 time bins leading up to (and including) time *t*: *X*_*t*_ ∈{0, 1}^7*N*^, where *N* = 93 is the number of neurons in the population. The evolution operator is then obtained by performing linear regression on Eq. 9, by minimizing the mean-squared residual 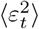. This gives 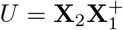 with the X_1_ =[X_1,_…,*X*_*T*-1_], X_2_ =[X_2,_…,*X*_*T*_] and 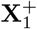 Moore-Penrose inverse of X_1_. To analyze the timescales of the dynamics and the corresponding features of the neural state, we perform an eigenmode decomposition of *U* : *Uv*_*i*_ = *λ*_*i*_*v*_*i*_ and 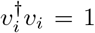. Writing the neural state in the eigenbasis, 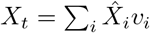, its evolution is given by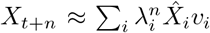. Thus, for each mode *v*_*i*_ and eigenvalue *λ*_*i*_, we can associate a timescale *τ*_*i*_ and frequency *ω*_*i*_ through:

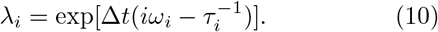

Note that the time scale *τ*_*i*_ diverges as |*λ*_*i*_|→1, which imposes |*λ*_*i*_ |≤ 1.

Next, we adapt the predictive IB described in the previous section to find the compression *Z*_*t*_ = Λ *X*_*t*_ that maximizes *I*_NCE_(*Z*_*t*_; *Y*_*t*_), where *X*_*t*_ is now the TDE of the neural state (of dimension 7*N*) re-defined in the previous paragraph, Λ is a *K* 7*N* matrix, and *Y*_*t*_ ∈{0, 1} ^*N*^ is the activity in a single time bin immediately following the last time bin of *X*_*t*_, as depicted in Fig. 4A. The difference from the IB described earlier is that *X*_*t*_ now contains the activity in the 7 time bins that lead to *t*. We made that choice to be able to make a connection with DMD, which predicts *X*_*t*+1_ as a function of *X*_*t*_, or equivalently *Y*_*t*_ as a function of *X*_*t*_, since the first 6 time bins of *X*_*t*+1_ are trivially the last 6 time bins of *X*_*t*_. In training, we choose *K* = 50 as this captures *>* 95% of predictive information.

**FIG. 4.**
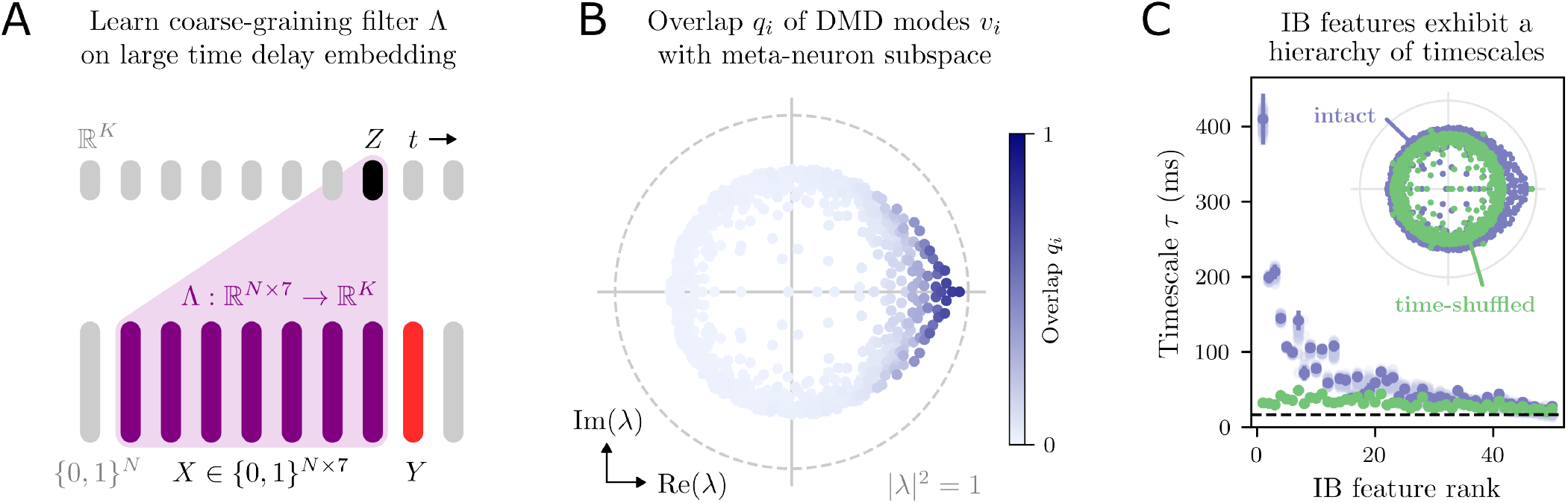
Meta-neuron features encode long timescales. All analyses were carried out on responses to opticflow stimulus. **A**. Compression of time-delay-embedded input variable *X*. We take a time delay embedding with 7 time steps, leading to a state *X* that is a 7*N* = 651-entry binary array. This is transformed linearly into *K* = 50 meta-neurons *Z*_*a*_ = Λ_*a*_*· X*, with *a* = 1, …, 50. That is, *Z* = Λ *· X*. **B**. Meta-neurons encode long-timescale DMD modes. Points are located at eigenvalues *λ*_*i*_ of the DMD operator *U*. Points closer to the unit circle indicate modes with longer timescales. Color depicts the overlap *q*_*i*_ defined as squared length of each DMD eigenmode *v*_*i*_ onto the vector space spanned by the Λ_*a*_ vectors. **C**. IB features are organized into a hierarchy of timescales. Performing PCA in the space of meta-neuron filters over 1000 repetitions of the training provides a hierarchy of IB features. Each feature *ϕ*_*µ*_ yields a timescale 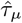 through projection onto the evolution operator, Eq. 12. Green points show the result for time-shuffled data that destroy predictability. Dotted line is at Δ*t* = 1*/*60 *s*. Error bars include uncertainty both in *U* and in *ϕ*. Light blue points show variation over 100 estimates of *ϕ*, each using only 500 repetition of training for Λ. Inset depicts the eigenvalues for *U* for both the intact (blue) and time shuffled (green) data.

In our first comparison between DMD and IB, we examine IB meta-neuron features in the eigenbasis{*v*_*i*_} of the DMD operator. Recall that IB meta-neuron features are encoded in the *K* row-vectors (each of dimension 7*N*) of the matrix Λ. Because any invertible transformation of *Z* leaves the mutual information invariant, all invertible linear operations Λ → *L*Λ yield equally good coarsegraining performance. What is important, then, is the rowspace of Λ, i.e. the linear subspace of ℝ^7*N*^ spanned by the row-vectors of the matrix Λ, which is invariant to Λ → *L*Λ transformations, and corresponds to what the meta-neurons collectively pay attention to in the neural activity *X*. Therefore, to quantify how well each mode *i* of the DMD dynamics is encoded in the IB meta-neurons, we define its overlap with their subspace as:

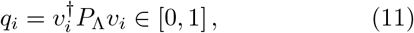

where *P*_Λ_ = Λ^*T*^ (ΛΛ^*T*^)^−1^Λ is the projection operator onto the rowspace of Λ. For example, if a mode has an overlap *q*_*i*_ = 1, then *X v*_*i*_ can be read out of the meta-neuron state *Z* with perfect fidelity, whereas an overlap of zero indicates that *Z* can tell us nothing about *X v*_*i*_. Fig 4B shows all eigenvalues *λ*_*i*_ in the complex plane, colored by their overlap *q*_*i*_ with the IB meta-neurons. We observe that modes with the largest overlap *q*_*i*_ have the largest |*λ*_*i*_|, corresponding to the longest time scales. This result suggests that the IB picks out features of the neural state that are the most stable over time, and thus have the largest predictive power for their own evolution. Due to the invariance of *Z* upon linear transformation, the above analysis does not allow us to directly pair individual IB meta-neuron features (rows of Λ) with DMD modes (*v*_*i*_). To obtain an approximate hierarchy of IB features and break the symmetry, we run 1000 independent trials of the IB training, and look for the main directions of variation of the IB meta-neurons. Explicitly, denoting by 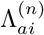the compression matrix in trial *n*, we compute its covariance 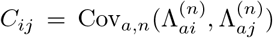 over both trial *n* and meta-neuron *a*, and diagonalize it: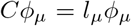 with 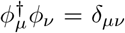. IB features are then defined by the eigenvectors *ϕ*_*µ*_, and ranked by descending order of explained variance *l*_*µ*_. We then express the evolution operator in this new eigenbasis: 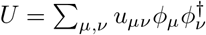. We observe that, empirically, diagonal terms *u*_*µµ*_ dominate that sum. Keeping only the diagonal terms and defining 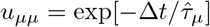, we obtain:

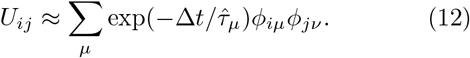

This rewriting of the evolution operator allows us to associate a time scale 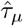 with each IB meta-neuron feature *ϕ*_*µ*_.

Fig. 4C shows that the time scale 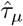 correlates with IB rank: it is highest for the most relevant IB features (highest *l*_*µ*_, low rank), and decreases as a function of rank (blue points). We emphasize that the IB features were not selected to exhibit this timescale hierarchy. As a control, we repeated the analysis on time-shuffled data (where both time and trial index are randomly permuted), revealing an almost flat dependence of time scale versus IB rank (green points), consistent with the expectation that time-shuffled data should have no predictive ability beyond the trivial 6-bin overlap between *X*_*t*_ and *X*_*t*+1_. Indeed, the DMD spectrum{*λ*_*i*_}of the intact versus time-shuffled data (inset) shows that large-modulus eigenvalues corresponding to long timescales are suppressed by this shuffling operation. In summary, this analysis shows that the features most relevant for prediction, as revealed by IB, correspond to the slowest modes of the dynamics revealed by the DMD analysis.

### F. Predictive information is collectively encoded

From the perspective of a downstream predictor, how important are collective effects in predicting retinal activity? To address this question, we first examine the contributions of individual neurons to the predictability of random groups. Specifically, we measure the relationship between a neuron’s information about itself (self-information) and the change in information it provides to a collective upon its inclusion (information contribution). For each neuron *A* in the 93-cell population, we compute its self-information *I*(*X*_*A*_; *Y*_*A*_) directly from an empirical histogram and sample ten random groups of 49 neurons *B* (Fig 5A). We then use NCE to estimate *I*(*X*_*B*_; *Y*_*B*_), the predictive information in these groups without *A*, as well as *I*(*X, Y*), the predictive information of all neurons in *A* ∪ *B* at the same interval *τ*. The difference δ_*A*|*B*_ = *I*(*X*; *Y*) − *I*(*X*_*B*_; *Y*_*B*_) represents the increase in total predictability upon including *A* in the collective. We call it the contextual information of neuron *A*.

**FIG. 5.**
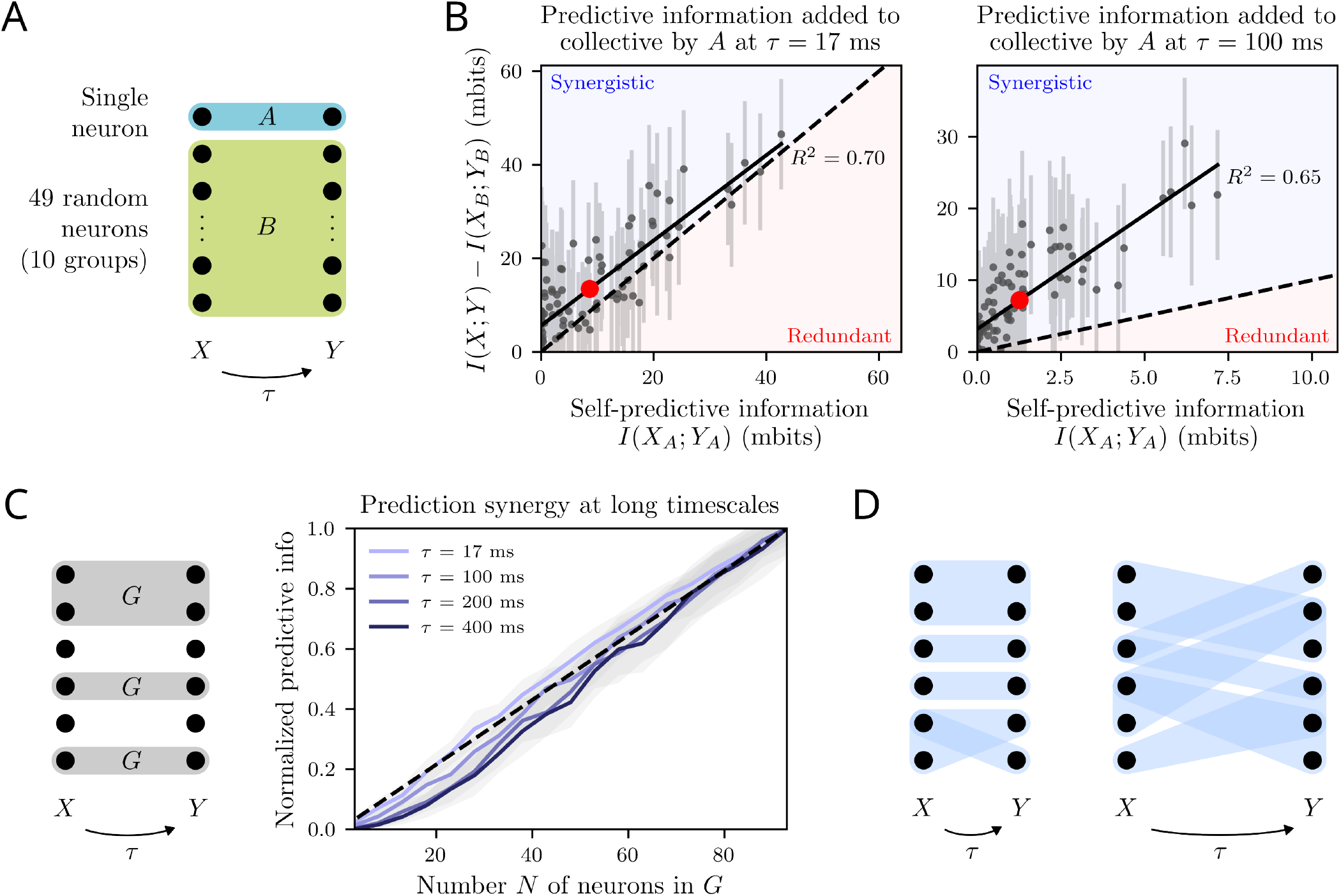
Synergy in predictive information. **A**. Depiction of single neuron contribution test. For each neuron *A* we randomly sample 10 groups *B* of 49 neurons each (*A* and *B* are disjoint) then estimate *I*(*X*_*A*_; *Y*_*A*_), *I*_NCE_(*X*_*B*_ ; *Y*_*B*_), and *I*_NCE_(*X*; *Y*), with *X* = *X*_*A*∪*B*_. We perform this test at *τ* = Δ*t* = 17 ms and at *τ* = 100 ms. **B**. Contextual predictive information δ_*A*|*B*_ = *I*(*X*; *Y*) − *I*(*X*_*B*_; *Y*_*B*_) of *A*, averaged over *B*, as a function of self-information *I*(*X*_*A*_, *Y*_*A*_). Deviation above the dashed unity line indicates synergestic effects. Red dot is the average over neurons. Collective effects are more important for prediction at the longer interval. **C**. Scaling of the mean predictive information 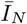 in 20 random groups *G* as a function of their size *N*, for four different prediction intervals. Each curve is normalized by its maximal value at *N* = 93. Curve convexity implies the presence of synergy, 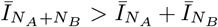. **D**. Schematic depiction of hypothesized correlation structure. At short timescales, prediction synergy is positive for small groups and neutral or redundant in large groups, suggesting greater autocorrelation content, less widespread cross-correlations, and effective independence between groups past some scale. Meanwhile, neurons which are highly self-predictive at long timescales also exhibit strong prediction synergy in random collectives, which is explained by strong cross-correlations.

Fig. 5B shows the contextual information, *δ*(*A B*), versus its self-information, *I*(*X*_*A*_; *Y*_*A*_), at short (17 ms) and long (100 ms) prediction timescales, for each neuron in the population. If neurons were encoding independent features of the stimulus, they would all fall on the identity line. We call neurons that encode more predictive information in the context of other cells than by themselves, *δ*_*A*|*B*_ *> I*(*X*_*A*_, *Y*_*A*_), synergistic. A vast majority of neurons are synergistic. In fact, many neurons have low self-information *I*(*X*_*A*_, *Y*_*A*_), but substantial contextual information *δ*_*A*|*B*_, especially at short *τ* = 17 ms (left panel). Such neurons are individually uninformative at the given timescale for downstream predictors; their utility only arises when read out within some group. These synergistic effects increase the overall predictability of the population: at short prediction internal (*τ* = 17 ms) the 93-cell population has about ⟨*I*(*X, Y*) ⟩ = 1.2 bits of predictive information per time bin, or about 13 mbits/neuron, versus *I*(*X*_*A*_; *Y*_*A*_) = 8 mbits of self-predictive information per neuron on average. Thus, correlations between weakly-firing, low self-information neurons and the other neurons cause the scaling of total predictive information to exceed the independent-neuron estimate. We find that synergistic effects are even stronger at longer timescales (*τ* = 100 ms, right panel), where predictive information is much lower, suggesting a shift towards collective encoding of predictive information as the prediction interval increases.

More generally, we can define a predictive synergy between two subpopulations *A* and *B* of neurons to quantify how much extra predictive information they carry together relative to their sum:

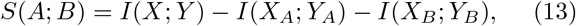

where *X* = *X*_*A*∪*B*_ is the joint activity of the population. To get an summary estimate of synergy in the population, we can compute the predictive information 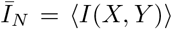as a function of *N*, averaged over random subgroups *G* of neurons of size *N*. Fig. 5C shows this quantity normalized by its maximal value at 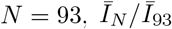, for various prediction intervals *τ*. We observe that 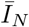is a convex function of *N*, which can be translated mathematically as 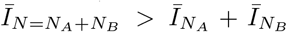, or equivalently *S*(*A*; *B*) *>* 0. The curve 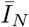 becomes more strongly convex, and hence predictive synergy larger, as the prediction interval Δ*t* is increased, consistent with the result for single neurons within a population.

We hypothesize that this trend towards increasingly collective encoding at longer timescales is reflected in the correlation structure, as schematically depicted in Fig. 5D, with blue connections representing correlations. A simple Gaussian toy model highlighting the relationship between correlations and synergy in presented in Appendix B. We explore more explicitly how correlations in the neural population impact synergy in the next section.

### G. How do correlations impact predictive information?

Correlations between neurons have two sources: stimulus-induced correlation, which stem from common or correlated stimulus inputs into the neurons, and noise correlations, which originate from shared noise and direct interactions between cells. To capture the contribution of noise correlations to predictability, we make use of the trial structure of the dataset (Figs 1E and 6A). Each stimulus was shown to the salamander retina around 80 times (in the case of the “opticflow” stimulus, 85 times). During each of these repeated trials, the stimulus driving responses in the retina stays exactly the same, so any variation in the responses comes from noisy outcomes of the internal dynamics of the retina. Formally, we can fix the stimulus by conditioning the response on the time *t* within the trial, *p*(*x, y*|*t*). Correlations between *X* and *Y* under this conditioning and across trials correspond to noise correlations. The unconditioned distribution studied in the previous section is then 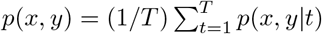, and includes both stimulus and noise correlations.

To remove noise correlations, we can randomly shuffle trials separately for each neuron. Denote by *x*_*n,t,a*_ the state of the *n*^th^ neuron on the *t*^th^ time step and in the *a*^th^ repeat of the stimulus. For each neuron, we draw a random permutation *π*_*n*_ that permutes trials to get a new dataset 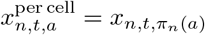 in which the noise correlation structure has been destroyed, while keeping the stimulus correlation structure through the time dependence. We call this operation, depicted on the left side of Fig. 6A, “per-cell” shuffle. It corresponds to making neurons conditionally independent: *p*_per cell_(*x, y t*) = ∏_*n*_ *p*(*x*_*n*_, *y*_*n*_ *t*), where the product runs over cells. Fig 6B shows that this per-cell shuffle does not have a substantial impact on predictive information at short timescales (*τ* = 1*/*60 s). This suggests that noise correlations between neurons do not contribute to the predictive information.

**FIG. 6.**
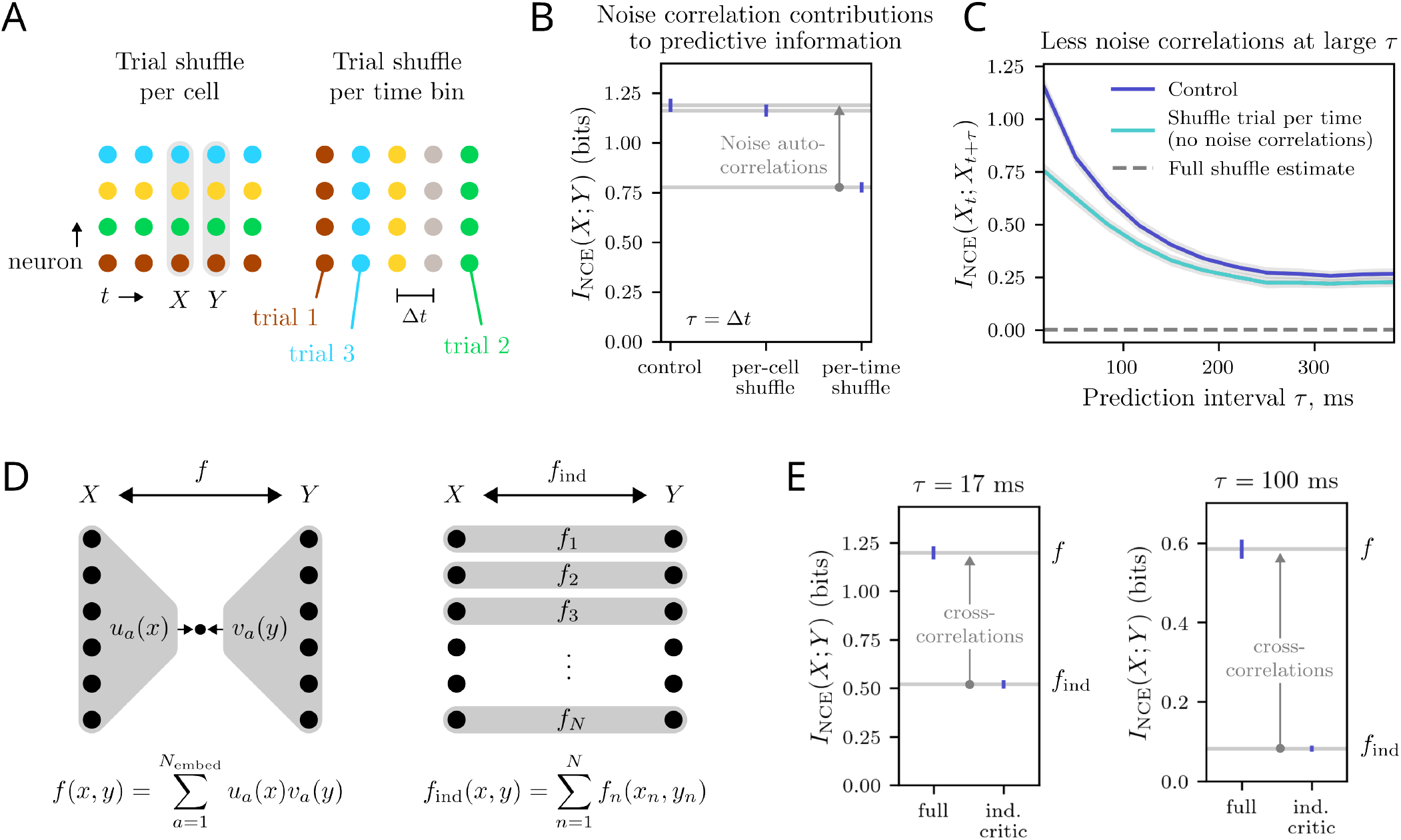
Contributions of cross- and autocorrelations to predictive information. Cross-correlations contribute significantly at short intervals and dominate at large intervals, while autocorrelations only significantly contribute at short intervals. Each stimulus (one of five 20 s naturalistic movies) was shown to the retina a total of 80-90 times. We use this trial structure to probe noise correlations. To remove noise correlations between neurons, we randomly permute the trial in a way that is consistent in time, but changes based on cell (left). We similarly remove noise auto-correlations by permuting the trial differently at each time step (right). Both these methods change the trial orderings but preserve the stimulus dependence. Predictive information from “opticflow” stimulus with different noise correlations removed by the methode described in A. Removing noise correlation between cells (per-cell shuffle) has negligible effect, but destroying noise autocorrelations (per-time shuffle) removes significant predictive information. **C**. Predictive information as a function of the prediction interval, with and without (per-time shuffle) noise correlations. At longer intervals, the gap narrows, meaning that noise autocorrelations matter less. **D**. By removing interactions between neurons in the critic function, we estimate the contributions of cross-correlations at all orders to predictive information. **E**. Difference in predictive information between fully expressive and independent critic functions, at two different prediction intervals. This difference represents the information discarded by a downstream predictor when it ignores cross correlations at all orders.

As a result of the negligible effect of noise crosscorrelations, any information carried from the present to the future by noise correlations must be comprised of autocorrelations, i.e. correlations between the activity *x*_*n*_ of a single neuron and its future *y*_*n*_, conditioned on *t*. We can quantify this effect by building a “per-time” shuffle, by drawing a trial permutation for each time *t, π*_*t*_, and defining shuffled data through 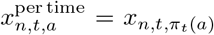. This other shuffling method, depicted on the right side of Fig. 6A, destroys correlations between the activities of neurons at different times conditioned on the stimulus, *p*_per time_(*x, y* |*t*) = *p*(*x*|*t*)*p*(*y*|*t*). Predictive information in that “per-time” shuffle is shown in Fig. 6B for *τ* = Δ*t* = 17 ms. Out of the 1.2 total bit of predictive information available at *τ* = Δ*t*, destroying temporal noise correlations removes around 0.35 bits. For longer prediction intervals *τ*, the relative contribution of noise correlations to predictive information begins to fade (Fig 6C). This tells us that the long timescales of predictions found in the neural code (quantified in the previous sections, see Fig. 2) are actually due to stimulus-induced correlations between neurons, rather than through the noise-induced auto-correlation of each neuron. In summary, noise correlations only contribute to predictive information at short intervals. Since these correlations are a single-neuron effect, their disappearance at longer timescales is consistent with our hypothesis that the balance of individual to collective correlations shifts towards collective at late times.

To quantify more directly the contribution to predictive information of correlations between neurons at different times, we need to go beyond shuffling operations. We would like to build a control in which correlations between different neurons at different times are destroyed, but not between neurons at the same time. We leverage the fact that the NCE estimate is a lower-bound of the Barber-Agakov bound; its optimization can be conceptualized as variational inference within a class of models. Recall that our variational estimate for a joint distribution *p*(*x, y*) is represented by a so-called critic function *f* (*x, y*), which encodes the interaction between *x* and *y*, but not the marginal joint distributions *p*(*x*) and *p*(*y*), which contain all same-time correlations between cells. So far we have taken the critic function to be a fully expressive neural network, able to capture any relevant correlations (Fig 6D, left). To destroy interactions between neurons across time, we consider an independent critic function ansatz (Fig 6D, right):

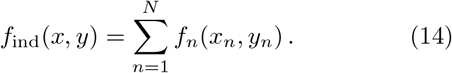

This independent critic yields a family of distributions *q*_ind_(*y x*) which only capture equal-time correlations and autocorrelations. In essence, this ansatz only allows each neuron to directly inform its own future.

Fig 6E shows the predictive information for the full (using a generic *f* (*x, y*)) versus independent (using *f*_ind_) critic for short (*τ* = 17 ms) and long (*τ* = 100 ms) prediction intervals, under the opticflow stimulus. The drop *D* in predictive information upon forcing independence may be written as:

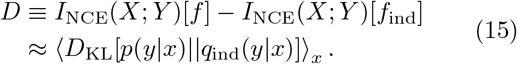

*D* represents the additional information about *Y* which can be extracted from *X* from temporal crosscorrelations. At *τ* = Δ*t*, observing cross-correlations provides about *D* = 0.7 bits, or 58% of the available predictive information, while at *τ* = 100 ms cross-correlations provide about *D* = 0.5 bits, or 84% of the total. This result directly confirms the picture that temporal crosscorrelations between neurons become increasingly important for prediction at long timescales.

## DISCUSSION

Combining information theory and machine learning we have demonstrated that the retina compresses information to make long-term predictions. By going beyond previous work that was limited to small sets of neurons [7, 44], we accessed the many-cell regime by leveraging a variational inference technique. We find that all predictive information in a large retinal population is encoded collectively in a few delocalized linear features. The predictive and decodable features we have discovered may be crucial for quick reactions in dangerous situations [45– 47].

In the large populations we studied, even once activity is binarized, the size of the space of possible states is 2^93^, suggesting that coarse-graining schemes are involved in neural processing. Additionally, since the neural code is sparse—neurons are much more likely to be silent— many states will never occur. Next, neural responses are correlated. These correlations are induced both by stimulus and by physiology, and both allow us to make statements about the state of given neurons given the firing patterns of other neurons. Finally, the code must ultimately be learnable, otherwise the brain could never decode visual signals.

A number of studies have investigated coarse-graining in the retina, and all describe procedures that reduce the state space while preserving relevant features [8, 24, 44, 48–52], although the definition of relevance varies between them. Here we focused on the predictive aspect and demonstrated a compression scheme which preserves predictive information in a relatively large population (93 RGCs). This allowed us to take large-scale collective effects into account and more accurately represent the problem presented to downstream neurons. In particular, we showed that predictive information in a population of retinal neurons can be encoded in a small number of linear features. By varying the target prediction interval, we found that compressibility extends to long timescales (500 ms), and that there are both generalized and timescale-specific predictive features present. Regardless of the timescale, these features are delocalized, spreading across the entire neural population, and capture information which is collectively encoded.

From the perspective of predictive downstream neurons, is it better to treat signals from neurons individually or collectively? We found that individual effects contribute most at short timescales, and collective effects are crucial at all intervals. We quantified the prediction synergy between different subsets of neurons. Single neurons almost always provide more predictive information to a collective than they have about themselves, and at moderate to long prediction intervals, we observed finite prediction synergy between large subsets. In order to see further into the future, downstream neurons therefore need to incorporate inputs from more neurons. Equal-time cell-cell noise correlations do not significantly contribute to predictive information. Noise autocorrelations are significant at short timescales, playing a smaller role for large prediction intervals. Even at the shortest prediction interval, close to half of the predictive information received by a downstream neuron comes from correlations between different RGCs. We conclude that although optimal prediction of retinal outputs at long timescales can be done with only a few linear features, these features need to be collective.

Our DMD analysis serves as a test which corroborates the idea put forward in [42], but in a complex biological context. By maximizing predictive information with a fixed set of variables, it seems we automatically encode long timescales. The correspondence suggests that if one chooses a number *K* of meta-neurons such that not *all* of the predictive information is captured, one still approximately recovers the *K* longest timescales in the system. To say with greater certainty that the *K* first IB features encode the *K* longest timescale modes, future work could try to find IB features which are true eigenfunctions of a better estimate of the DMD operator. This might be possible with more expressive, nonlinear meta-neuron features, as well as DMD with a well-chosen set of nonlinear observables.

By variationally solving a predictive information bottleneck problem on data taken from vertebrate retina, we have demonstrated that downstream prediction of a large population RGCs is plausible. Applied to data from these downstream neurons, future analyses can reveal their decoding performance relative to the optimum. Further work should constrain the coarse-graining maps and critic function Ansatz to a family of interpretable, mechanistic models of retina and downstream neurons and reveal how predictive computation might be enabled or hindered by physiology, similarly to [49]. More generally, given the ubiquitous importance of prediction as a biological function, we also anticipate that this method can be useful in finding computationally relevant coarse-graining schemes in other complex and biological systems.

## DATA AND CODE AVAILABILITY

Data is available at https://doi.org/10.5061/dryad.4qrfj6qm8. Code is available at https://github.com/sepalmer/metaneurons.

## ACKNOWLEDGMENTS

This work was supported by the National Science Foundation through the Physics Frontier Center for Living Systems (PHY-2317138). This work was supported by the NSF-Simons National Institute for Theory and Mathematics in Biology, awards NSF DMS-2235451. This was work supported by a FACCTS award (AMW, SEP). This work was partially supported by the European Research Council Consolidator grant no. 724208 (AMW), Fondation Bettencourt-Schueller (TM), Simons Foundation MP-TMPS-00005320, as well as a Schmidt Sciences Polymath award (SEP).

## Appendix A NCE implementation details

Here we give details on how the “critic function” is computed and optimized. Recall that it takes the form:

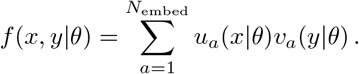

We take *v*_*a*_ and *u*_*a*_ to be feed-forward multilayer perceptron (MLP) neural networks and *θ* are all of the model parameters. When compressing from a single time bin, both *u* and *v* have 2 hidden layers with 32 neurons, and output to the embedding dimension of size *N*_embed_ = 30. When we compress from a TDE state of seven time bins, we use hidden layers with 32 neurons and an embedding dimension of 100. Hidden layers have a tanh nonlinearity. The input dimensions depend on the specific calculation. For example, when coarse-graining to *K* = 4 meta-neurons to predict the whole population state in a single future time bin, we would have *u* : ℝ^4^ → ℝ^30^ and *v* : ℝ^93^ → ℝ^30^.

To train the models, we used the Adam optimizer [53, 54] with a learning rate of .009, batch sizes of *B* = 800, and held out half the data for testing. All reported mutual information estimates are from the held-out test sets. We explored effects of dropout on weights within these MLPs but found that early stopping with no dropout was the most effective regularization. We chose to stop training after each data point in the test set had been used by the optimizer 16 times. In both testing and training, the first 20 frames (1/3 of a second) from each response set were removed as these contain transients from stimulus switching. For testing, we constructed 500 batches of size *B* = 500. Typically, this whole train/test procedure would be done on the order of 10 s of times, producing means across batches, training initializations, and random train/test designations. Uncertainty for mutual information estimates incorporated both noise due to different initializations as well as noise between batches

## Appendix B Prediction synergy

To understand how correlation structure may affect prediction synergy, we briefly consider a Gaussian toy model. Two sub-systems *A* and *B* have “present” states *x* = (*x*_*A*_, *x*_*B*_), and future states (*y*_*A*_, *y*_*B*_). Because all variables are all jointly Gaussian, we can specify all parameters by choosing first- and second-order correlation functions. We choose a ll meanstobe zero, that is for *α* = *A, B*, ⟨*x*_*α*_⟩ = 0 = ⟨*y*_*α*_⟩. Next, we fix the scale of fluctuations so that all 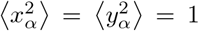. With only pairwise correlations between degrees of freedom left, we choose three parameters, *a, b*, and *c*, representing autocorrelations, equal-time *A*-*B* correlations, and across-time *A*-*B* correlations (cross-correlations) respectively. Note that the only choices of (*a, b, c*) which are valid are those such that all correlation matrices are positive definite. The expression for prediction synergy in this toy model is

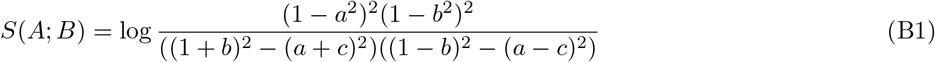

As a point of reference, note that the predictive information of either subsystem is given by

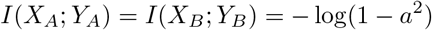

Where for ⟨ [*x*_*A*_, *y*_*A*_]^*T*^ [*x*_*A*_, *y*_*A*_] ⟩ to be positive definite we must have *a*^2^ < 1.

Several limits of Eq. B1 are especially illuminating. The easiest limit is that of independence between the subsystems, *b* = *c* = 0. While this can be seen to yield *S*(*A*; *B*) = 0 from the expression above, the deeper reason is that when *A* and *B* are statistically independent we must have *I*(*X*; *Y*) = *I*(*X*_*A*_; *Y*_*A*_) + *I*(*X*_*B*_; *Y*_*B*_), and hence *S*(*A*; *B*) = 0.

Next, consider a model where *B* is a direct copy of *A*. Since *x_A_* has the same relationship with *y*_*A*_ as it does with *y*_*B*_, we must have *a* = *c*. Further, *x*_*A*_ must relate to *x*_*B*_ in the same way it relates to itself, so we take *b* → 1, which yields *S*(*A*; *B*) = log(1 − *a*^2^) = − *I*(*X*_*A*_; *Y*_*A*_). That is, the prediction synergy is negative, indicating redundancy. Moreover, the extent of this redundancy is given by the total predictive information of a single subsystem, in agreement with our interpretation of this model as consisting of two redundant copies.

As a final limit, we consider the removal of all correlations except cross-correlations. In this case, *a* = 0 implies that *I*(*x*_*A*_; *y*_*A*_) = 0, meaning each subsystem observed alone has no information about its future. Taking *b* = 0 as well, we find that *S*(*A*; *B*) = − 2 log(1 − *c*^2^). From the definition of *S*, we also see that *S*(*A*; *B*) = *I*(*X*; *Y*). In this limit, all predictive information is encoded collectively, in that observation of both subsystems is required to extract it. It is interesting to note that the expression − 2 log(1− *c*^2^) is the same as in the case of independent systems except under the replacement *c*→ *a*. The basic reason for this is that this limit also describes two independent subsystems, but *x*_*A*_ should be grouped with *y*_*B*_ and *x*_*B*_ with *y*_*A*_, meaning our choice of subsystems does not capture this independence.

